# Context dependent function of the transcriptional regulator Rap1 in gene silencing and activation in *Saccharomyces cerevisiae*

**DOI:** 10.1101/2023.05.08.539937

**Authors:** Eliana R Bondra, Jasper Rine

**Author notes:** For correspondence: Jasper Rine.

## Abstract

In *Saccharomyces cerevisiae,* heterochromatin is formed through interactions between site-specific DNA-binding factors, including the transcriptional activator Rap1, and Sir proteins. Despite a vast understanding of the establishment and maintenance of Sir-silenced chromatin, the mechanism of gene silencing by Sir proteins has remained a mystery. Utilizing high resolution chromatin immunoprecipitation, we found that Rap1, the native activator of the bi-directional *HML*α promoter, bound its recognition sequence in silenced chromatin and its binding was enhanced by the presence of Sir proteins. In contrast to prior results, various components of transcription machinery were not able to access *HML*α in the silenced state. These findings disproved the long-standing model of indiscriminate steric occlusion by Sir proteins and led to investigation of the transcriptional activator Rap1 in Sir-silenced chromatin. Using a highly sensitive assay that monitors loss-of-silencing events, we identified a novel role for promoter-bound Rap1 in the maintenance of silent chromatin through interactions with the Sir complex. We also found that promoter-bound Rap1 activated *HML*α when in an expressed state, and aided in the transition from transcription initiation to elongation. Highlighting the importance of epigenetic context in transcription factor function, these results point toward a model in which the duality of Rap1 function was mediated by local chromatin environment rather than binding-site availability.

**Significance Statement:** The coarse partitioning of the genome into regions of active euchromatin and repressed heterochromatin is an important, and conserved, level gene expression regulation in eukaryotes. Repressor Activator Protein (Rap1) is a transcription factor that promotes the activation of genes when recruited to promoters, and aids in the establishment of heterochromatin through interactions with silencer elements. Here, we investigate the role of Rap1 when bound to a promoter in silent chromatin and dissect the context-specific epigenetic cues that regulate the dual properties of this transcription factor. Together, our data highlight the importance of protein-protein interactions and local chromatin state on transcription factor function.

## Introduction

Cellular identity can be defined by the array of expressed and repressed genes in a cell. Thus, two cells with identical genomes can exhibit vastly different phenotypes. Due to the wide variety of expression patterns needed for normal development and function, eukaryotic gene expression is controlled by many different processes ranging from gene-specific combinatorial effects of transcription factors to domain-wide compaction or accessibility of chromatin. Further, modifications to chromatin promote differential regulation via both the recruitment and restriction of transcriptional activators and repressors (1). The coarse partitioning of the genome into regions of actively expressed euchromatin and repressed heterochromatin is a characteristic of eukaryotic genomes and a major point of gene expression regulation (2). The stability of cell type is controlled, in large part, by the faithful propagation of cell-type-specific patterns of gene expression over cellular divisions. Breakdown of finely tuned expression programs can lead to aberrant gene expression, disease, or cell death.

In *Saccharomyces cerevisiae*, heterochromatin is controlled by the Silent Information Regulator (Sir) proteins which assemble at the cryptic mating-type loci, *HML* and *HMR,* and the telomeres (3–5). Study of the recruitment and spread of these proteins has been fundamental in understanding the establishment and maintenance of heterochromatin (6). The canonical view of the establishment of silencing posits that Sir proteins are recruited to nucleation sites termed silencers (7–10). The *E* and *I* silencers, negative *cis-*regulatory sequences, flank both *HML* and *HMR* and are the sites from which Sir proteins spread across these loci in a sequence-independent manner. Recent evidence from our laboratory indicates that, in addition to these silencers, the promoter of *HML (HML-p)* acts as an early nucleation site of silencing (11). Common to all three of these early-recruitment loci (*HML-E, HML-I,* and *HML-p)* is the presence of a binding site for Repressor activator protein 1 (Rap1) (8, 10, 12, 13).

Rap1 is best characterized in its role as an essential transcription factor that activates hundreds of genes across the genome including the majority of ribosomal protein genes (14–18). Much of Rap1 research has focused on the activator function of the protein. *In vitro* studies of Rap1 classify it as a “pioneer factor”, a term used to described a class of proteins that are unique in their ability to bind to DNA in the presence of nucleosomes, establish domains of open chromatin, and facilitate binding and recruitment of other transcription factors (18–22). In addition to the Sir proteins, Rap1 functions in establishing and maintaining silent chromatin at *HML, HMR,* and telomeres by binding to silencers and recruiting Sir proteins, in combination with two other silencer-binding proteins, the Origin Recognition Complex (ORC) and the transcription factor ARS-binding factor 1 (Abf1) (23–29). How Rap1 mediates two apparently opposing functions has remained a mystery.

Despite decades of research utilizing Sir silenced chromatin as a model for heterochromatic gene repression, the fundamental question regarding the mechanism of Sir-based silencing has remained inadequately answered. In the most broad-scale model, Sir proteins form a macromolecular complex that blocks, wholesale, protein-DNA interactions in silent chromatin, including transcription factors accessing their cognate binding sites. This model is supported by evidence of expression state-dependent cleavage and modification of enzyme recognition sites in silent or active chromatin (30, 31). A more nuanced version of this mechanism supports specific pre-initiation complex interference by Sir proteins. Here, DNA-binding activators access their binding sites in Sir-silenced chromatin, but subsequent assembly of a functional pre-initiation complex is somehow hindered (32, 33). Yet other work suggests a downstream-inhibition model whereby silencing acts by prohibiting formation of mature transcripts rather than transcriptional initiation, and is based on results indicating no difference in recruitment of TATA-Binding Protein (TBP) nor RNA Pol II to silent chromatin, but instead a marked absence of elongation factors and mRNA capping machinery (34, 35). Thus, the extent to which transcription machinery is occluded, and the specificity of such blockage, has remained inconclusive.

Sequence identity between the mating-type locus *MAT* and the auxiliary *HML* allows a unique opportunity to study the role of Rap1 in both silent and expressed contexts, and how local chromatin state affects function. The promoters of *MATα* and *HMLα* are identical in sequence and, thus, each contains a Rap1 binding site. Rap1 binding at *MAT* is responsible for activation of *α1* and *α2* (36, 37). The presence of this same promoter binding site at *HML*, which is constitutively silenced, offers an opportunity to test predictions of the various models of silencing. While it is generally understood that Rap1 binding at the silencers *HML-E* and *HML-I* recruits Sir proteins to mediate silencing, the role for Rap1 at the promoter is posited to be an activator (8, 36, 37). To date, it is unclear to what extent Rap1 binds its recognition site in a heterochromatinized context, and whether it contributes to either silencing or activation of the *HML* locus.

Prior studies have been unable to query the endogenous activator at the *HML* due to the difficulty of distinguishing binding at this locus to binding at *MAT*. To better understand the dichotomy of Rap1 function, we utilized endogenous tagging of the protein in combination with high-resolution ChIP-seq and RNA measurements to characterize the contributions of Rap1 to silencing and expression at *HML*. We investigated the *in vivo* residence times of Rap1 to further characterize the interaction between Rap1 and chromatin.

## Results

### Rap1 bound to the promoter of *HML* in a silenced state but failed to recruit transcription machinery

Previous studies addressing the mechanism of silencing have yielded contradictory and sometimes paradoxical results, due in part to the low-resolution techniques used at the time (32, 34, 35, 38, 33). In studies attempting to characterize the limiting steps of Pol II recruitment to a silenced locus, regions of sequence identity between *MAT, HML* and *HMR* have interfered with the unambiguous assignment of recruitment of different factors to the loci (figure 1A). To ensure unambiguous interpretation of our results regarding recruitment to *HML*, we designed and performed experiments in strains lacking *MATα* and wherein the *α2* coding sequence at *HML* was replaced by the coding sequence for the *Cre* recombinase (*hmlα*2Δ::*Cre*), thus avoiding confounding sequence identity. Similarly, to characterize enrichment at *MATα*, we performed experiments in strains lacking both *HML and HMR (hml*Δ *hmr*Δ).

**Figure 1.**
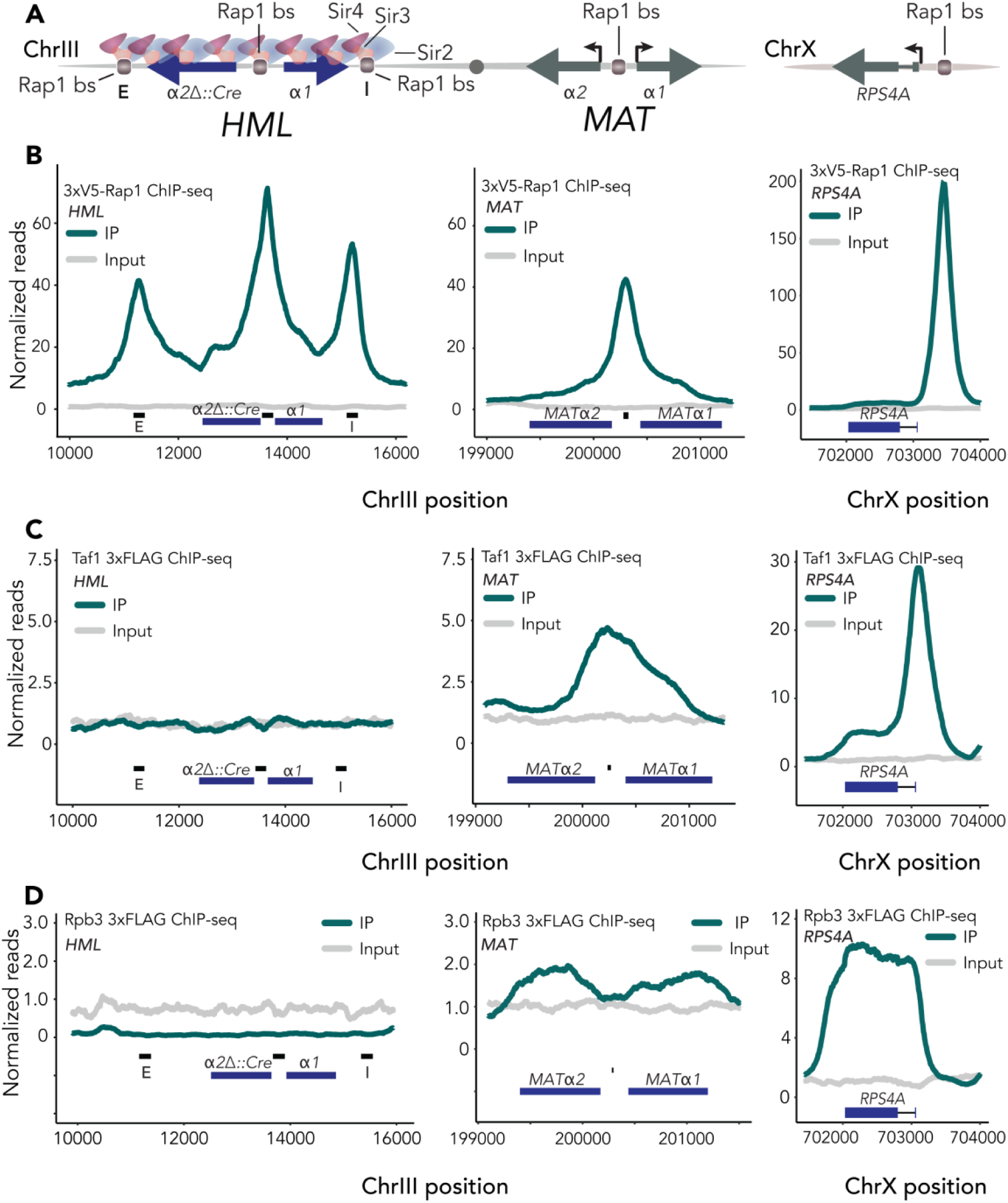
Rap1 bound the promoter of *HML* in a silenced state but failed to recruit the pre-initiation complex. For all ChIP-seq experiments, read counts were normalized to the non-heterochromatic genome-wide median. IP and input values are plotted on the same scale. Data shown are the average of two ChIP-seq experiments unless otherwise noted. (A) Schematic of *HMLα* and *MATα* on chromosome III. Rap1 binding sites at *HML-E*, *HML-I,* and the promoter of both *HML* and *MAT* are noted. (B) Left, averaged normalized reads for ChIP-seq in two 3xV5-Rap1 samples at *HML* in *SIR* cells. Black bars represent 200 bp surrounding Rap1 binding sites at *HML-E, HML-p* and *HML-I,* respectively. Middle, same as left but showing *MAT.* Right, same as (left,middle) but at *RPS4A*. (C) Same as (B) but for Taf1-3xFLAG-KanMX. (D) Same as (B,C) but for Rpb3-3xFLAG-KanMX.

We tagged endogenous Rap1 with 3xV5 at the N-terminus in order to retain both its essential activating and repression functions, permitting accurate representation of Rap1 enrichment at *HML* and *MAT* (figure S1A,B). Utilizing Chromatin Immunoprecipitation followed by next-generation sequencing (ChIP-seq) we determined that Rap1 was in fact bound to the promoter of *HML* in silenced chromatin (figure 1B). Compared to the extent of enrichment at *MATp*, Rap1 was unexpectedly enriched at the silenced locus relative to the unsilenced (figure 1B).

Strong enrichment of Rap1 at the *HML* promoter under wild-type conditions was incompatible with the generalized steric-hindrance model and led us to reconsider the remaining hypotheses; either that silencing occurs at some point after the recruitment of trans-activators but before that of RNA Pol II, or silencing blocks elongation, analogous to paused RNA polymerase II in other eukaryotes (32, 34, 35, 38, 33). To distinguish between these mechanisms, we endogenously tagged a set of proteins intimately involved in RNA Pol II-dependent transcription: TATA binding protein-Associated Factor 1 (Taf1), RNA Polymerase B 3 (Rpb3), and Elongation Factor 1 (Elf1). As with our tagged Rap1, epitope tags did not affect fitness of cells with the tagged versions as the only forms of these proteins in the cell (figure S1A,B).

TFIID is one of the first factors recruited to transcription initiation sites (39–41). A subunit of TFIID, Taf1, has proposed interactions with Rap1 making it a compelling protein of interest for assessing recruitment of transcription machinery to silent chromatin (42, 43). In contrast to previous reports, TFIID showed no enrichment at *HML* in silenced chromatin (figure 1C). It was therefore unsurprising that neither a major subunit of RNA Pol II, Rpb3, nor the elongation factor Elf1 exhibited any binding to silenced chromatin (figure 1D, figure S1C). As an internal positive control, we mapped enrichment of each protein at *MAT* in *hmr*Δ *hml*Δ cells, where the recruitment of each followed expected patterns; the initiation factor Taf1 was localized over the promoter, while the RNA Pol II subunit Rpb3 was enriched over the gene bodies (figure 1C,D). Furthermore, all three proteins were substantially enriched at *RPS4A,* a ribosomal protein gene that is also a known Rap1 target (figure 1, figure S1C). These data revealed that Sir-silenced chromatin was not entirely refractory to protein binding, but specifically to RNA pol II transcription machinery. In sum, we found robust recruitment of the endogenous activator to native Sir-silenced *HML* and narrowed the step at which silencing occurs to a point between recruitment of the activator, Rap1, and the formation of the pre-initiation complex.

### Rap1 contributed to the maintenance of silent chromatin at the native *HML* promoter

Given that Rap1 was enriched at the *HML* promoter in silenced chromatin, but TFIID was not, we investigated the possibility that promoter-bound Rap1 contributed to silencing the locus. We generated a strain with a two base-pair mutation in the Rap1 binding site at the promoter which is known to strongly decrease expression of *α1* αnd *α2* (36, 37). Upon mutating GG to TC, we saw significant reduction of Rap1 at its consensus binding sequence within the *HML* promoter (figure 2A, figure S2C,D). Introduction of this binding site mutation did not affect enrichment of Rap1 at other loci genome-wide (figure S2A).

**Figure 2.**
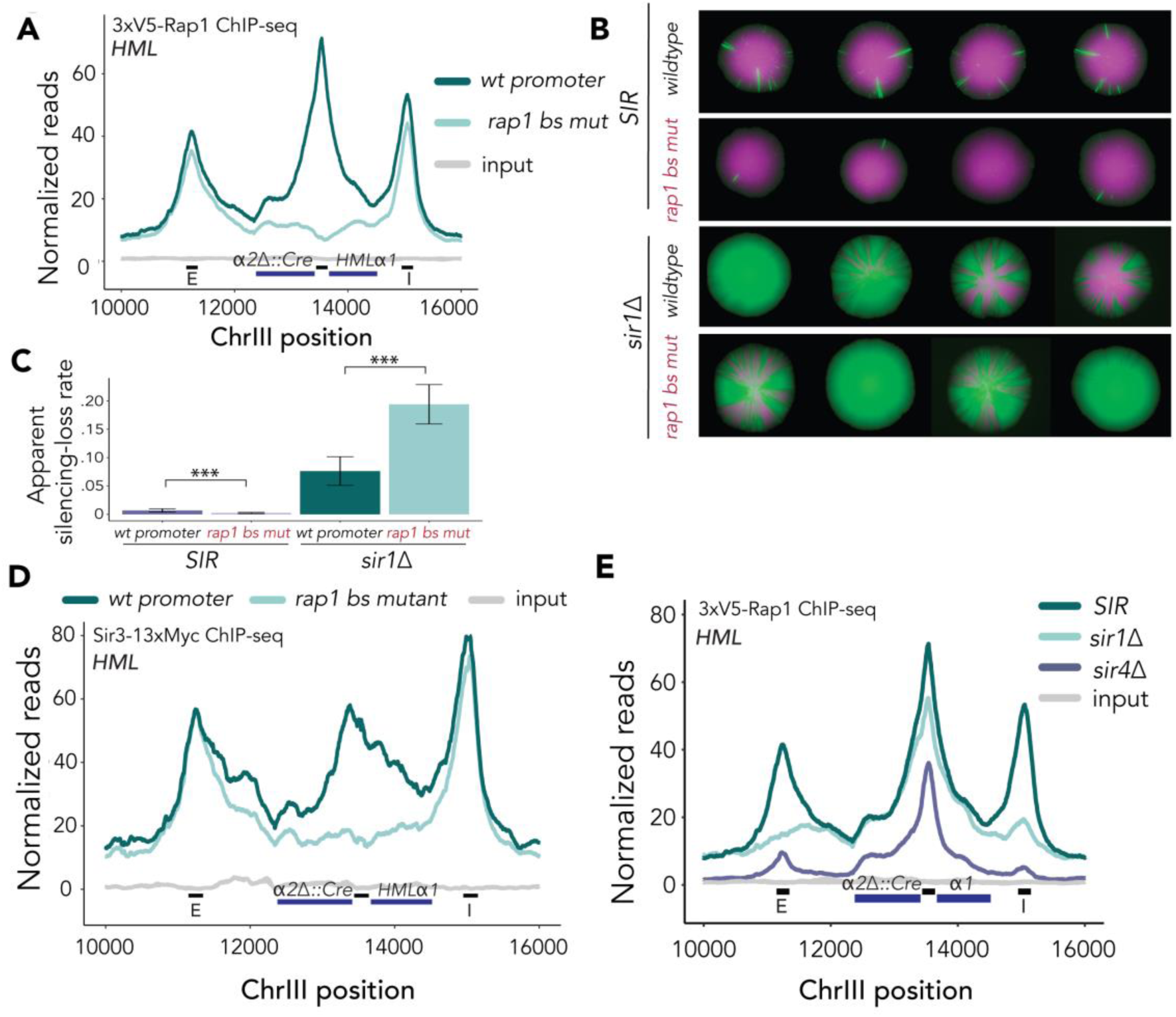
Rap1 contributes to the maintenance of silent chromatin at the native *HML* promoter Unless otherwise stated, ChIP-seq data represented averaged reads of two biological replicates over the locus, normalized as in figure 1. Black bars along x-axis represent 200 bp surrounding Rap1 binding sites at *HML-E, HML-p,* and *HML-I,* respectively. IP and input values are plotted on the same scale. (A) Normalized reads mapped to *HML* in two 3xV5-Rap1 ChIP-seq experiments for wild-type and mutant Rap1 binding motif at the promoter. (B) Representative CRASH colonies for *SIR* and sir1Δ cells with wild-type and mutant Rap1 binding site at *HML-p*. (C) Apparent silencing-loss rate for genotypes described in (B) ± SD. The following number of events was recorded for each sample: *SIR* wt promoter (n = 271933); *SIR* rap1 bs mutant (n = 773105); *sir1*Δ wt promoter (n = 151846); *sir1*Δ rap1 bs mutant (n = 90211). p-values (p<2.2e-16) for both comparisons were calculated using a two-sided t-test. (D) Normalized ChIP-seq reads for Sir3-13xMyc mapped to *HML* for wild-type and mutant Rap1 binding motif at the promoter. (E) Normalized ChIP-seq reads for 3xV5-Rap1 mapped to *HML* in *sir1*Δ, *sir4*Δ and *SIR* cells.

To evaluate the impact of the Rap1 binding site mutation (rap1 bs mutant) on silencing at *HML*, we introduced the two base-pair mutation into a previously developed strain that monitors loss-of-silencing events (44). This assay, Cre-Reported Altered States of Heterochromatin (CRASH), allows for highly sensitive measurements of loss of silencing events by expression of *HMLα2Δ::Cre* and a subsequent recombination event that results in a unidirectional switch from red fluorescence to green fluorescence (figure 2B top panel, figure S2B) (44). Silencing is a robust process which fails approximately once in every 1000 cell divisions (44, 45). To increase the level of expression to a measurable amount and broaden the dynamic range, we deleted the *SIR1* gene (*sir1*Δ) in a strain with the rap1 bs mutation and the CRASH background. *sir1*Δ cells exist in a bimodal state of expression at *HML* (46). Recent evidence has shown that even silenced *sir1*Δ cells exhibit reduced binding of all other Sir proteins across the locus, thereby representing a weakened heterochromatic domain (47). Interestingly, mutation of the Rap1 binding site at the *HML-*promoter in *sir1*Δ cells did not show reduced sectoring (figure 2B, bottom panel). We utilized flow cytometry to quantify changes to the silenced domain observed with the CRASH assay (48). Surprisingly, in *sir1*Δ cells the apparent silencing-loss rate was higher in rap1 bs mutant cells than in those with the wild-type promoter (figure 2C), indicating that promoter-bound Rap1 contributed to silencing at *HML*.

We hypothesized that promoter-bound Rap1’s intrinsic enhancement of silencing may be through strengthening the stability of silent chromatin. To assess this, we performed ChIP-seq of a Myc-tagged allele of Sir3 as a proxy for enrichment of the Sir complex across the locus. Congruous with our finding that apparent silencing-loss rate was higher in the weakened Sir state of *sir1*Δ cells, Sir3 occupancy was reduced in rap1 bs mutant cells (figure 2D). This reduction was particularly striking over the promoter, showing an approximate 3-fold reduction in Sir3 occupancy at this locus. In contrast, Sir3 enrichment at *HML-E* and *HML-I* was unaffected. Although Sir3 enrichment was reduced by deletion of the Rap1 binding site, measurements in Sir-competent cells by both CRASH (figre 2B) and RT-qPCR (figure 3A) revealed that cells were able to maintain silencing. This further supported a model in which interactions between Sir proteins and Rap1 cooperate to form and maintain silenced chromatin. Together, these experiments showed that Rap1 bound to the same locus could perform opposing functions dependent on nuances in the local chromatin environment.

**Figure 3.**
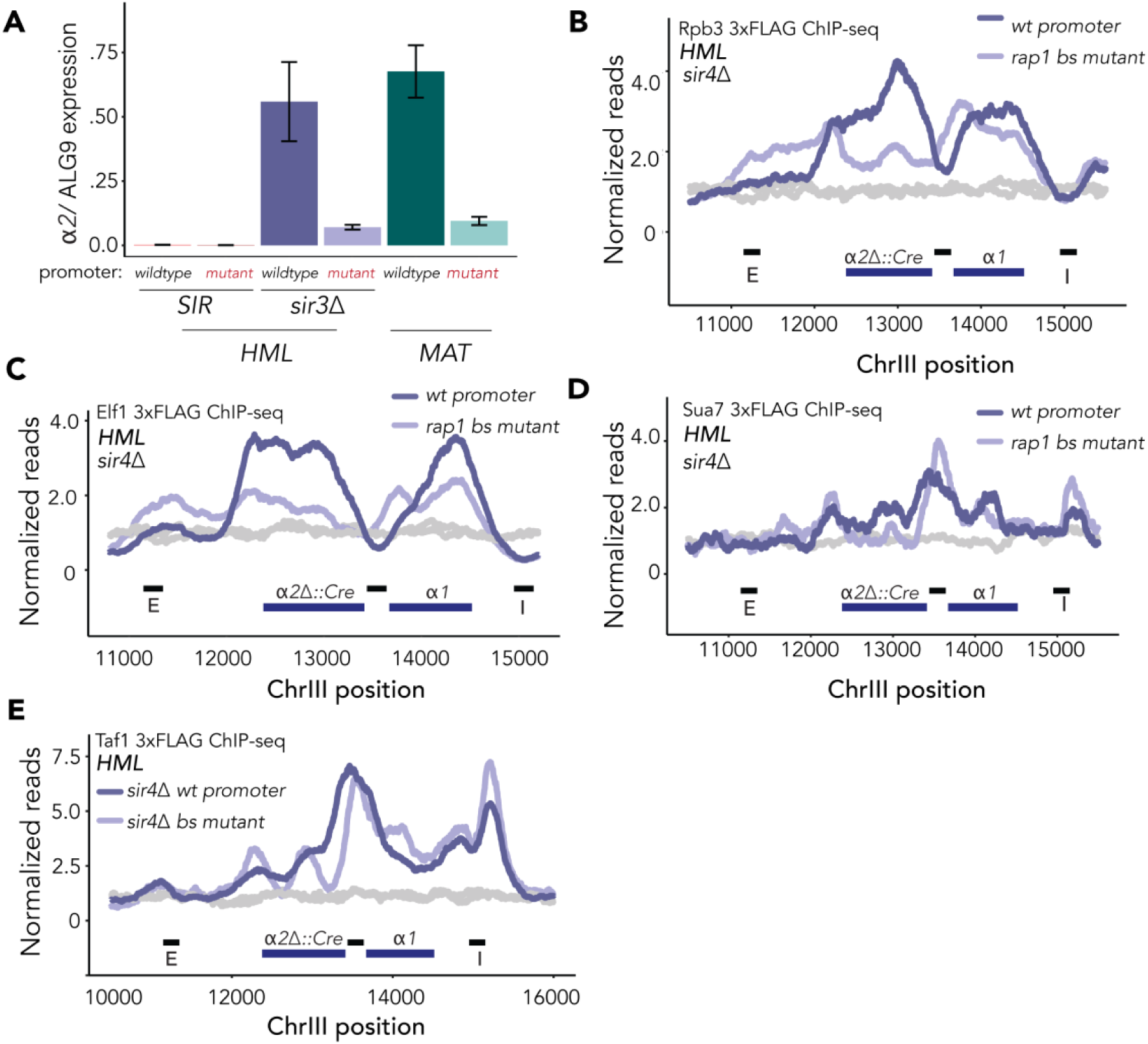
Promoter-bound Rap1 was able to activate transcription of unsilenced *α2* and aided the transition from initiation to elongation at unsilenced *HML*. For all ChIP-seq experiments, read counts were normalized to the non-heterochromatic genome-wide median. IP and input values are plotted on the same scale. Data shown are the average of two ChIP-seq experiments, unless otherwise noted. (A) RT-qPCR quantification of *α2* expression at *HML* and *MAT* normalized to control locus *ALG9*. Each plot consists of an average of 2 biological replicates with 3 technical replicates for each. Error bars represent ± SD. (B) Normalized reads for ChIP-seq of Rpb3-3xFLAG at *HML* in *sir4*Δ cells. (C) Same as (B) but for Elf1-3xFLAG-KanMX. (D) Same as (B,C) but for Sua7-3xFLAG-KanMX. This dataset is from only one sample. (E) Same as (B,C) but for Taf1-3xFLAG-KanMX.

Cooperativity between Sir proteins and Rap1 would predict that enrichment of Rap1 at the promoter may be decreased by the absence of Sir proteins. We therefore performed Rap1 ChIP-seq in *sir4*Δ cells, in which the Sir complex is absent from the *HM* loci (figure 2E). Rap1 enrichment at *HML-p* in silenced cells was found to be significantly greater than that in unsilenced cells (figure 2E, S2D,H; Student’s t-test p = 0.024). We also found a substantial decrease in Rap1 occupancy at the silencers *HML-E*, *HML-I,* and *HMR-E,* which has a Rap1 binding site important for silencing, despite sequence-specific recruitment of Rap1 to these loci preceding Sir protein recruitment in canonical models for the establishment of silencing (figure 2E, S2D, figure S2G). To test whether diminished occupancy of Rap1 at *HML* in *sir4*Δ cells was due to a disruption of the interaction between Sir4 and the C-terminal domain of Rap1, we performed ChIP-seq in *sir3*Δ cells and found the results to be nearly identical (figure S2E,F). Again, as an internal positive control, Rap1 enrichment at *MATα* was found to be similar to that at unsilenced *HML* and less than the enrichment at *HML* in silenced chromatin (figure 1B, 2E, S2H). As expected, Rap1 binding at *MATa* did not vary based on the availability of the Sir complex (figure S2G).

To expand upon the finding that Rap1 enrichment varied with local availability of Sir proteins, we performed ChIP-seq of Rap1 in *sir1*Δ cells (figure 2E), acknowledging that the ratio of cells with silenced or expressed *HML* loci differ between cultures. Rap1 was enriched to an intermediate level at *HML-p* in these cells (figure 2E). Furthermore, we found a relative decrease in Rap1 at the silencers at the silencers where Sir1 is known to play a critical role in silencing establishment and has a direct effect (figure 2E) (10, 14, 38, 49, 50). Collectively, these data inferred that cooperative interactions existed between Sir proteins and Rap1 (figure 1B, figure 2E). Taken together these findings established a novel and specific contribution of promoter-bound Rap1 to silencing, where it was previously thought to have potential only for activation.

### Promoter-bound Rap1 activated transcription of unsilenced *α2* and aided the transition from initiation to elongation at unsilenced *HML*

Rap1 is required for transcription of *α2* and *α1* at *MAT* (37). Therefore, by abrogating Rap1 enrichment at the cognate *HML* promoter site we presumably disrupted expression of the locus to some degree. To assess the extent to which Rap1 at the *HML* promoter had the potential to also serve as a transcriptional activator at this site, we performed reverse-transcription quantitative polymerase chain reaction (RT-qPCR) to measure mRNA expression from the *HMLα* locus in Sir+ and Sir-cells. Expression of *HMLα* was nearly undetectable in *SIR+* cells. However, in *sir4*Δ cells, which have no Sir protein recruitment to the locus, expression of *α2* from *HML* was comparable to its expression from *MAT* (figure 3A). In contrast, we saw an approximate 5-fold reduction in *HMLα2* expression in rap1 bs mutant cells as compared to wild-type *HML-p,* which was comparable to the reduction caused by the same binding site mutation at *MAT* (figure 3A). This result was consistent with our finding that rap1 bs mutant cells have lower rates of sectoring than their wild-type promoter counterparts in the CRASH assay (figure 2B, second panel). These data confirmed previous work on the role of Rap1 at *MAT* (36, 37), and extended those conclusions by establishing that Rap1 contributes significantly and equivalently to expression of *α1* and *α2* at both the unsilenced *HML* and native *MAT* loci (figure 3A).

To understand which step of transcription Rap1 contributed to the most, we performed ChIP-seq of tagged proteins in *sir4*Δ cells with and without the rap1 bs mutation at the promoter. As predicted by the decrease in gene expression (figure 3A), enrichment of major Pol II subunit Rpb3 and elongation factor Elf1 over the gene bodies of *HMLα2* and *α1* was decreased (figure 3B,C). We noted, however, a non-canonical binding pattern of these proteins over the bi-directional promoter. Rather than exhibiting the same decreased occupancy as the coding sequence, Elf1 and Rpb3 were enriched at the promoter in rap1 bs mutant cells relative to their wild-type counterparts (figure 3B,C). This pattern is indicative of a failure in promoter escape, or the transition to productive elongation (51). Furthermore, we found enrichment of TFIIB subunit Suppressor of AUG 7 (Sua7) over the promoter in rap1 bs mutant cells relative to wild type cells (figure 3D). TFIIB typically dissociates from the promoter at the initiation stage and does not travel with RNA Pol II as it transcribes (52). Together these findings demonstrated a role for Rap1 in promoter escape of actively transcribed genes.

### *In vivo* Rap1 residence time did not correlate with differences in function at *HML* and MAT

ChIP-seq offers a static view of protein-DNA interactions across the genome. In light of recent focus on protein dynamics as a critical lens through which to study transcription, we hypothesized that the dynamics of Rap1–DNA interactions may vary between heterochromatin and euchromatin, due to the distinct compositions of the two structures. To test this hypothesis, we utilized the rapid nuclear depletion strategy afforded by the anchor-away technique to remove unbound Rap1 from the nucleus (figure S3A) (53). Due to the nature of the anchor-away experiment, wherein Rap1 was depleted over time, it was important to include a spike-in control for downstream analysis of the ChIP-seq data. We used cells from the closely related species *Saccharomyces paradoxus* which allowed unique mapping of sequences from each species (54, 55). Normalizing to number of reads in each sample assigned to *S. paradoxus,* we fit ChIP-seq enrichment data to a non-linear regression model as described by the DIVORSEQ method (56). This allowed us to calculate the apparent k_off_ for each peak, and thus a proxy for the *in vivo* residence time (figure S3A). We characterized the fits and apparent residence times for 377 Rap1-bound loci across the genome, in replicate, at which Rap1 binding decayed over time (figure S3B-G). Finally, we generated a series of synonymous single nucleotide polymorphisms to *HML* to allow unambiguous assignment of high-throughput sequencing reads to either *MAT* or *HML* within the same sample (figure 4A) (57). The residence time of Rap1 at the promoter in silent (*HML)* and active *(MAT)* chromatin was similar, although initial Rap1 enrichment was decreased at *MAT* as seen previously (figure 4B). The dwell-time of Rap1 bound to silencers was also similar (figure 4C,D). These results indicated that the dual functions of Rap1 could not be attributed to differences in dynamics, but rather resulted from local chromatin contexts and possibly other protein-protein interactions.

**Figure 4:**
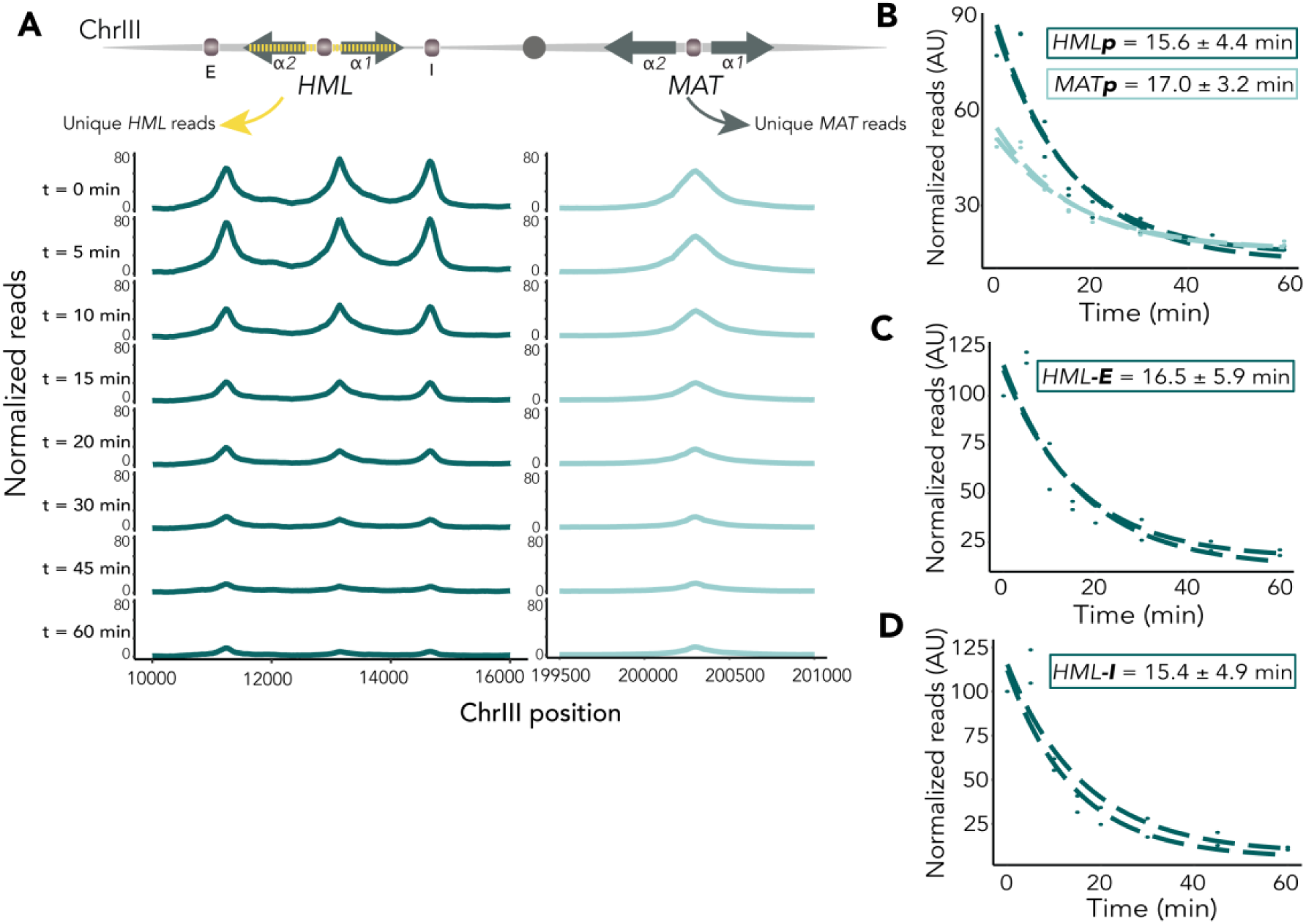
*In vivo* Rap1 residence time did not reflect differences in chromatin state Decay of Rap1 occupancy at *HML* and *MAT* by anchor away (A) Top, schematic of introduction of SNPs to enable unique mapping of *HML* and *MAT* in a strain that contains both. Below, Rap1 enrichment by two, averaged ChIP-seq experiments at *HML* (left) and *MAT* (right) over time-course, plotted on the same y-axes. (B) Fitted non-linear regressions for residence times of *HML-p* and *MAT-p.* Each replicate is shown separately. ± SE of average residence time (C) Fitted non-linear regressions for residence times of *HML-E* as in (B). (D) Fitted non-linear regressions for residence times of *HML-I* as in (B), (C).

### Genome-wide analysis of *in vivo* Rap1 apparent residence times

As chromatin state did not appear to contribute to differences in Rap1 apparent off-rate, we tested whether other context-specific cues contributed to this metric. Using cutoffs similar to those previously described (56), we refined a set of 1118 Rap1-bound regions genome-wide and ultimately measured Rap1 off-rate at 377 of these sites (figure 5A, figure S4). To assess the contribution of Rap1 apparent dwell-time to function in transcriptional regulation, we selected Rap1 enrichment peaks that were within 500 bp upstream of open reading frame and assigned peaks to these respective genes. This dataset was then subdivided into quartiles based on residence time: shortest (n = 95), short (n = 95), long (n = 93), longest (n = 94). Of note, 42 of the 96 subtelomeric peaks (defined as located within 15 kb from the ends of telomeres) displayed poor fits due to a lack of decay over time (figure S4C). These peaks were almost exclusively the Rap1-bound loci at the most telomere-proximal positions, or within 500 bp of the ends of chromosomes. Conversely, subtelomeric peaks that ranged from 500 bp – 15 kb from the ends of telomeres exhibited shorter dwell-times relative to telomeric regions, though, ultimately, dwell-time did not correlate with distance from chromosome end (figure S4E,F). These findings underscored a difference in dynamics between Rap1 bound at the very ends of chromosomes, which presumably functions as a structural element in telomere end-protection, and Rap1 bound at subtelomeric loci, which may be involved in heterochromatin-mediated silencing.

**Figure 5.**
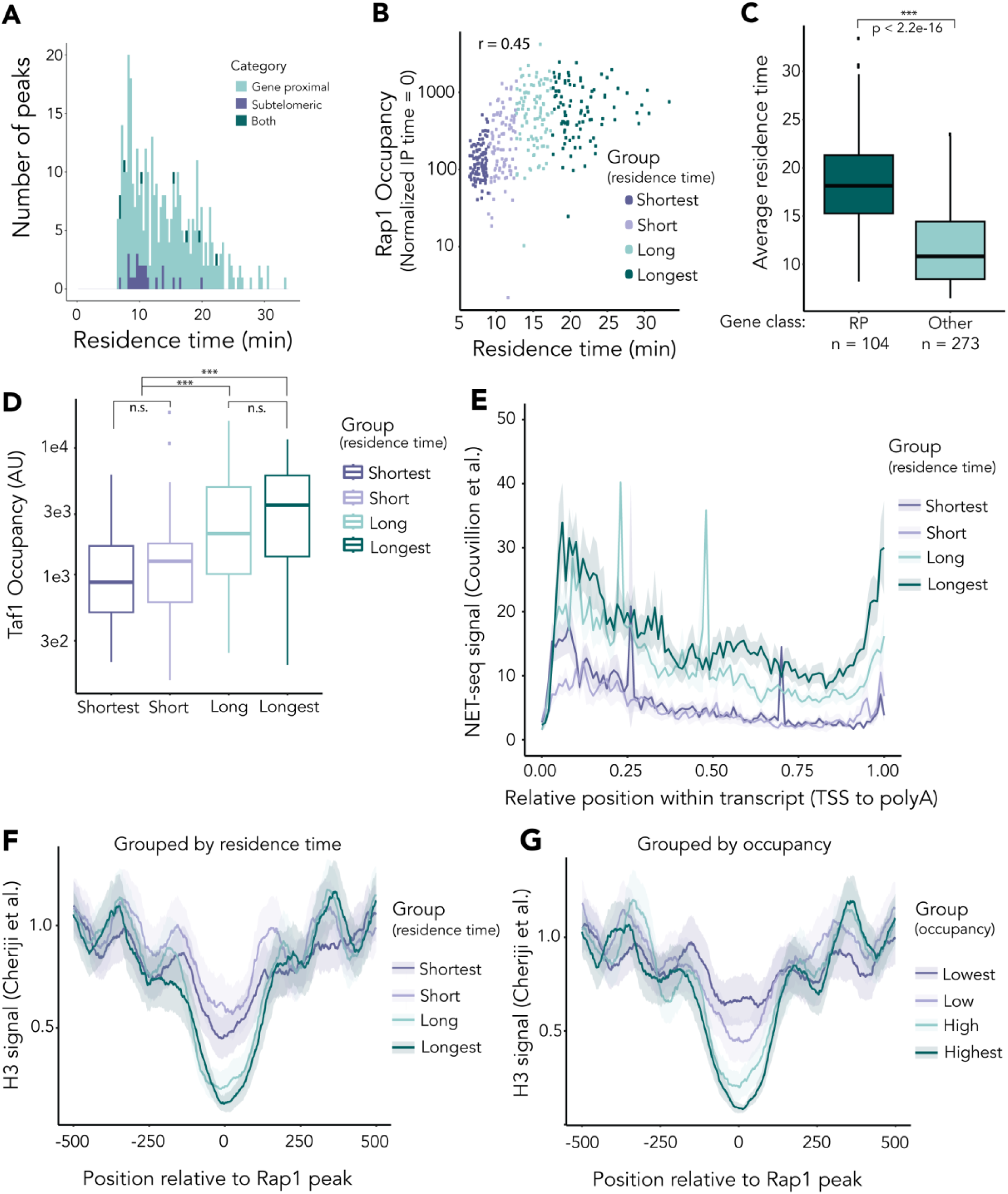
Genome-wide analysis of *in vivo* Rap1 apparent residence times supports and extends previous models that Rap1 dwell-time is correlated with transcriptional output All figures comprise data obtained from the average of two biological replicates. Peak set (n=377) was divided into quartiles based on residence time for analysis unless otherwise noted. For gene-level analyses, Rap1 peaks were assigned to ORFs for which a Rap1 peak summit was within 300bp upstream of ORF start. (A) Average apparent Rap1 residence time in minutes of the 377 binding sites evaluated genome-wide categorized as gene-proximal, subtelomeric, or both. (B) Correlation between average Rap1 occupancy before depletion (enrichment at t=0) and the average apparent residence times for all 377 Rap1-bound peaks (Pearson correlation *r* = 0.45, p-value < 2.2e-16). (C) Difference in apparent residence times between sites that are classified as regulating Ribosomal Protein genes (n = 104, dark green) and all other sites (n = 273, light green). The p-value was calculated using a Mann-Whitney U test (p < 2.2e-16). (D) Quantification, by mean apparent residence time quartile, of normalized Taf1 occupancy levels at the Rap1 binding sites. These levels are defined as the amount of Taf1 enrichment (in reads) covering the Rap1-bound loci. Significance was calculated using a one-way ANOVA followed by Tukey’s HSD test (p < 2.2e-16). (E) Mean profiles display NET-seq coverage (61) with 95% confidence intervals (displayed as transparent filling) within neighboring transcript(s). Coverage was scaled according to transcript length. (F) Summary distribution plots of average H3 enrichment (64) centered on Rap1 peaks and spanning 500bp+/-. Coverage was grouped by apparent-residence time quartiles. The confidence intervals are indicated as in figure 5E, with a transparent fill denoting 95% confidence intervals. (G) Same as F, but peaks grouped by ranked Rap1 occupancy at t=0.

As ChIP-seq peak heights reflect a static view of occupancy of chromatin binding proteins, we tested the correlation between Rap1 occupancy (enrichment at time=0) and apparent dwell-time (figure 5B). There was a significant positive correlation between these two factors (Pearson correlation coefficient *r* = 0.45, p-value < 2.2e-16), but much of the variation remained unaccounted for. Rap1 targets vary greatly in expression. Indeed, ribosomal protein genes whose transcripts make up nearly 60% of total mRNAs in the yeast cell (58) were enriched for longer apparent dwell-times as reported previously (figure 5C) (59). Presumably, stable Rap1 binding would allow for efficient recruitment of pre-initiation complex machinery via TFIID–Rap1 interactions (43). Utilizing our previously generated dataset of Taf1 ChIP-seq as a proxy for TFIID occupancy, we compared dwell-time to Taf1 enrichment. Overall, there was a positive correlation between apparent residence time and Taf1 ChIP signal (figure S4G, Spearman correlation coefficient ρ = 0.43, p-value < 2.2e-16), and a significant difference in Taf1 occupancy between the shorter and longer Rap1 dwell times (figure 5D, ANOVA, p < 2.2e-16). To investigate further the role of Rap1 binding dynamics in transcription regulation, we generated summary distribution plots (meta-gene analyses) of nascent transcripts associated with the Rap1-bound loci genome-wide. Utilizing published datasets (60, 61), we plotted nascent, elongating Pol II occupancy reported by Native Elongating Transcript sequencing (NET-seq) for each transcript proximal to a Rap1 peak. This meta-gene analysis showed that longer apparent Rap1 dwell-time correlated with greater NET-seq signal and thus an inferred higher transcriptional output (figure 5E). These data reinforced the proposed model in which longer apparent dwell-time corresponded to higher transcriptional activity, perhaps working through stable recruitment of the pre-initiation complex.

Many studies identify Rap1 as an important modulator of nucleosome-free regions through interactions with the chromatin remodeling complex RSC (18, 19, 62, 63). *In vitro*, introduction of nucleosomes to naked DNA is anti-correlated with stable Rap1 binding (21). We therefore hypothesized that, similarly to other transcription factors, apparent Rap1 dwell-time would be inversely correlated with nucleosome occupancy, and that Rap1 binding stability was related to its role in determining nucleosome-free regions. Using a published dataset that utilized a chemical cleavage method to precisely map single nucleosomes with high accuracy genome-wide in yeast (64), we created a meta-gene analysis mapping histone H3 positioned 500 bp upstream and downstream of each Rap1 peak (figure 5F). The averaged nucleosome occupancy for each quartile group of residence times revealed that longer dwell time corresponded to a broader nucleosome-free region, with a notably weakened −1 positioned nucleosome (figure 5F). We then compared how nucleosome occupancy corresponded to Rap1 occupancy (IP enrichment at time = 0). When positioning data were grouped by initial enrichment instead of dwell-time, we found a consistent anti-correlation between histone occupancy centered over the Rap1 peak and relative Rap1 occupancy (figure 5G). As total enrichment is a function of both on-rate and off-rate, we surmised that the differences we observed in correlations between nucleosome occupancy and Rap1 enrichment versus apparent dwell-time may have reflected differences in on-rate. In sum, these data supported and extended the hypothesis that Rap1 binding and nucleosome occupancy were inversely correlated *in vivo*, and connected this attribute to transcriptional output.

## Discussion

This study explored the enigmatic ability of Rap1 protein to function in both gene activation and repression. Earlier work discounted the possibility that the function of Rap1 was determined by subtle differences in its recognition sequence (65). Other work suggested that Sir-silenced chromatin could inhibit binding of at least some proteins to their recognition sites (32, 34, 35, 38). In contrast, our findings established that silenced chromatin did not block Rap1’s recognition of its binding site in silent chromatin. Instead, our data revealed a more nuanced and unexpected way in which Rap1 and Sir proteins mutually reinforced each other in the assembly of silenced chromatin.

Rap1, bound to its recognition sequence within the bi-directional promoter between *HML α1* and *HMLα2* served opposing functions depending on the state of the surrounding chromatin. The *HML* promoter binding site was robustly enriched for Rap1 in the presence of Sir proteins. In contrast, transcription initiation factors were undetectable. Strikingly, Rap1 at the bi-directional promoter internal to *HML* resulted in increased enrichment of Sir proteins at *HML*, and vice versa. Moreover, *HML-*promoter-bound Rap1 contributed to the stability of silent chromatin when weakened (figure 2). In addition, we determined a role for Rap1 in promoting the transition from initiation to elongation, which may be dependent on its role in nucleosome positioning.

Our results agreed with earlier work that Sir-silenced chromatin can inhibit some proteins from binding their recognition sequences (32, 33, 35, 38). Despite ample Rap1 enrichment at a Sir-silenced *HML,* we found no evidence of the pre-initiation complex bound at the *HML* promoter in silent chromatin, nor any indication that silencing acts by blocking Pol II elongation (figure 1). Our study used tagged forms of Taf1, Sua7, Rpb3, and Elf1, each expressed from their native promoter as proxies for TFIID, TFIIB, RNA Pol II, and elongation machinery, respectively. Previous reports find TATA-binding protein (TBP) and TFIIH to be present in the context of Sir-silenced chromatin (35). The earlier study utilized ChIP in conjunction with PCR and did not provide the resolution of the current study. Although we did not evaluate TFIIH subunits, TFIIH recruitment to a locus follows, and is dependent on, TFIID binding (66), and we did not find evidence of TFIID enrichment in the presence of Sir proteins (figure 1). Relatedly, we chose Taf1 as the proxy for the pre-initiation complex based on studies revealing nearly all genes in the yeast genome are TFIID-dependent and that Taf1 is necessary for recruitment of TBP and assembly of the pre-initiation complex (PIC) (67–69). In summary, our results demonstrate that Sir-silencing occurred through a pre-initiation complex interference mechanism, whereby the presence of Sir proteins competitively inhibited the ability of Rap1 to recruit the transcription machinery.

While apparent off-rates for Rap1 did not differ between silenced and unsilenced chromatin, we found greater enrichment of Rap1 in the presence of Sir proteins both at the promoter and silencers of *HML* (figures 1,2,4). Taken together, we inferred that the enhanced ChIP-based enrichment of Rap1 in the presence of Sir proteins was consistent with an increase in the frequency of Rap1 binding events in this context. However, inferences regarding off-rate through the anchor-away method are limited by the method affecting primarily the loss of free Rap1, and any changes in on-rate are speculative. Nevertheless, our dwell-time data were consistent with those described by a related but different technique that reports on local competition between Rap1 molecules (59). Our data supported a hypothesis that interactions between Sir proteins and Rap1 resulted in greater recruitment of both to silenced chromatin than would be achieved by the affinity of Rap1 for its binding site alone.

The *sir1*Δ genotype has long been used as a case study in epigenetics, as *sir1*Δ cells exist in a bi-stable population of silenced and unsilenced *HM-*loci despite having the same genotype (46, 70). Our samples of *sir1*Δ cells were prepared from unsorted cultures. The *sir1*Δ-*rap1* bs mutant strain revealed two distinct patterns of Rap1 enrichment across *HML* (figure S2C). Notably, one sample almost exactly matched the enrichment pattern seen in a *sir4*Δ-*rap1* bs mutant. From this we inferred a higher rate of cell-switching to a more *sir4*Δ-like state when Rap1 was absent from the *HML* promoter. Furthermore, we found the apparent silencing-loss rate in *sir1*Δ-*rap1* bs mutant cells to be greater than in a *sir1*Δ alone (figure 2B,C). In the *sir1*Δ mutant, the density of Sir proteins at the silent locus is less than in wild type, creating a paucity of Rap1-Sir interactions and, more broadly, a decrease in stability of the silent locus (47). Combined, these data reflected the importance in Rap1-Sir protein interactions particularly over the promoter in the weakened *sir1*Δ silent domain. The weakened interactions could allow for more opportunities for transcription machinery to interact with Rap1 and shift the balance in favor of transient derepression. We propose that Rap1 acts as a toggle in the competition between transcription and silencing of *HML*.

By indirect means, the activation domain of Rap1 has been shown to interact with various TFIID components (43, 71). Despite the activation domain and C-terminal interaction domain appearing to be non-overlapping by those analyses, one explanation for the occlusion of the pre-initiation complex from Sir-silenced chromatin would be if binding of Sir proteins to Rap1 rendered the activation domain inaccessible to TFIID. Our data were compatible with the idea that interactions between Rap1 and Sir proteins are mutually exclusive to interactions between Rap1 and TFIID subunits. In this model the balance of silencing versus activation would tip slowly toward fully silenced as the local concentration of Sir proteins increased.

In agreement with previous findings of other transcription factors, our data showed that nucleosome positioning was correlated with Rap1 dwell-time (56). In our anchor-away dataset, Rap1-bound loci with the longest apparent residence time were characterized by a well-defined nucleosome-free region centered on the Rap1 binding site, and a broader nucleosome-depleted region with an unstable −1 positioned nucleosome (figure 5F). Conversely, loci with shorter dwell-times corresponded with a narrowing of the nucleosome-free region centered on the Rap1 peak, and an associated relative enrichment in the presence of histones over these peaks (figure 5F). These data suggested that Rap1 binding in the presence of a nucleosome was less stable, as is reported *in vitro* (21). We identified a further anti-correlation between nucleosome occupancy and Rap1 occupancy (enrichment at t=0, figure 5G). By taking both the static occupancy measurement and the apparent dwell-time function into account, we inferred a variable on-rate that supports this hypothesis. These data indicated that nucleosome positioning relative to Rap1 peak summit played an important role in determining both the level to which Rap1 was enriched at those sites, and the apparent residence times once bound. However, it would be equally valid to infer that strong Rap1 binding depleted nucleosomes to a greater extent than weak Rap1 binding.

Based upon the positive correlation between Rap1 residence time and enrichment of Taf1, we hypothesized that the variability in expression strength at different Rap1-bound loci may be due in part to variability in Rap1 dwell-time (figure 5D, S4G). Furthermore, nascent transcript abundance correlated with Rap1 dwell-time (figure 5E). Thus, we have confirmed and extended previous models suggesting the role of Rap1 in transcriptional activation was dependent, in part, on Rap1 binding dynamics (19, 62, 63, 72). In summary, apparent residence time may be attributed to competition with nucleosomes, with the creation of a stable nucleosome depleted region allowing for higher rates of transcription.

Rap1 occupancy is an important determinant of the size and patterning of the nucleosome depleted regions to which it binds, while removal of Rap1 may result in remodeling of the nearby chromatin (18, 62, 63,73–75). Sir protein avidity for nucleosomes creates a robust pattern of nucleosome occupancy at *HML* and *HMR* (76). Alteration to the nucleosome depleted region in the bi-directional promoter in the absence of Rap1 could generate a block to productive transcription. We observed a narrowed Taf1 peak in rap1 bs mutant cells, which may indicate reduced access of TBP to the TATA-box (figure 3E). Rap1 binding is necessary for downstream recruitment of the chromatin remodeler complex RSC to maintain a nucleosome depleted region (19). Without Rap1 binding, and recruitment of chromatin remodelers, these sites may be less accessible to subunits of TFIID and other pre-initiation complex machinery. Furthermore, in the context of unsilenced *HML,* Elf1 and Rpb3 appeared to pile up over the bi-directional promoter in the absence of Rap1, which indicated a blockade in the progression of transcription (figure 3). Our finding that the switch from RNA Pol II initiation to elongation was hindered in the absence of Rap1 could reflect a change in the promoter architecture upon removal of Rap1.

Rap1 has been described as a pioneer factor with the ability to access cognate binding sites in the presence of a nucleosome array *in vitro* (21, 77). Rap1 bound to its promoter site in Sir-silenced chromatin extends this view *in vivo* (figure 1). In a broader context, pioneer factors are typically utilized by the cell during periods of drastic genomic restructuring, such as during fertilization in metazoans. Their pervasive binding to regions of the genome allows for poising of the genome for activation of cell-fate-specific gene expression (78, 22). In their haploid life cycles, yeast continuously undergo chromatin landscape restructuring in the form of DNA replication during replicative aging. Furthermore, it is important for the single-celled organism to readily adapt to environmental stresses. Rap1 is necessary for the Gcn4-mediated regulation of ribosomal protein genes which occurs upon amino acid starvation (79). Perhaps the downstream effect of Rap1-mediated nucleosome-free regions is to more readily enable the genome to activate certain genes under stress conditions by promoting the transition from transcription initiation to elongation.

In summary, we found that Rap1 has a complex and context-dependent role in the regulation of gene expression, with the ability to both stabilize the Sir-silencing complex at a silenced promoter and promote transcriptional elongation at the same locus in the absence of Sir proteins. In this way, Rap1 can be compared to Glucocorticoid receptor, a transcription factor well studied for its context-specific roles in vertebrate gene regulation (reviewed in 88, 89), and Ume6, a meiotic regulator that can act as a repressor or activator depending on its cofactors (82). Like the glucocorticoid receptor, one possible explanation of how Rap1 may be able to bind DNA in heterochromatin but not recruit transcription machinery may be the presence of post-translational modifications to the Rap1 protein. In a thematically similar concept, recent data reveals differential phosphorylation of Clr4^SUV39H^ correlates with a switch in methylation state of H3K9 in *S. pombe* (83, 84).

This work highlights the many modes of epigenetic regulatory mechanisms integrated by cells to give rise to a vast spectrum of context-specific and finely tuned gene expression patterns. Beyond the role of Rap1 in *S. cerevisiae,* these findings have implications, broadly, in eukaryotic regulation of cell-type fidelity across cell divisions. Dual-function transcription factors can be recruited to promoters and, in a context-dependent manner, serve as co-activators or co-repressors to finely tune gene expression, in part through the effects of local concentration of interaction partners. In conclusion, these findings provide new insights into the mechanisms of gene expression and highlight the importance of considering the context in which transcription factors function.

## Acknowledgments

We are grateful to the Rine laboratory for helpful discussions in the planning of this work. We give special thanks to Davis Goodnight for his experimental guidance and keen editing, and to Marc Fouet for his generosity with all things microscopy. We thank Paige Diamond for her invaluable assistance with data analysis and discussion. We also thank Elçin Ünal and Danielle Hamm for providing critical feedback on this manuscript. This work relied on the Vincent J Coates Genomics Sequencing Laboratory at UC Berkeley. This work was funded by grants from the National Institutes of Health to JR (R35GM139488). EB received support from a National Science Foundation Graduate Research Fellowship (Grant No. 1752814) and NIH Training Grant (T32GM007232).

## Author Contributions

Eliana Bondra: Conceptualization, Data curation, Formal analysis, Validation, Investigation, Methodology, Writing - original draft, review and editing.

Jasper Rine: Conceptualization, Supervision, Funding acquisition, Writing - review and editing.

## Data Availability

All ChIP-seq datasets (raw and processed) are available at NCBI Gene Expression Omnibus (GEO): Series GSE227763.

## Materials and Methods

### Yeast strains

Strains used in this study are listed in *SI Appendix, Dataset S1*. All strains were derived from the *S. cerevisiae* W303 background (except JRy15212 which was derived from *S. paradoxus YPS138*) using standard genetic techniques and CRISPR-Cas9 technology (85–87). Deletions were generated using one-step replacement with marker cassettes (88, 89). Details of strain construction for epitope-tagged proteins and mutants can be found in *SI Appendix, Supplementary Methods.* Relevant oligonucleotides used for strain construction can be found in *SI Appendix, Dataset S2*.

### CRASH colony imaging

Colonies were plated onto 1.5% agar plates containing yeast nitrogen base without amino acids, 2% dextrose, and supplemented with complete supplement mixture (CSM)-Trp to minimize background fluorescence. Colonies were incubated for 5–7 days at 30 °C, then imaged as described in (90).

### Flow cytometry and calculations of apparent loss-of-silencing rate in CRASH strains

This experiment was carried out as described in (Janke et al 2018 and Fouet and Rine 2023). To summarize: strains were streaked for single colonies on YPD, with multiple single colonies used as technical replicates for each sample. Strains were then back-diluted in growth medium containing G418 to select for cells that had not yet lost silencing. Cells were diluted and grown in liquid CSM until mid-log phase and harvested by centrifugation, then resuspended in PBS at approximately 0.5 OD. Samples were processed as described in Fouet and Rine 2023. The apparent silencing-loss rate was calculated as previously described (48, 90) (*SI appendix, Supplementary Methods*).

### RNA extraction and RT-qPCR

RNA extraction and RT-qPCR was carried out as in (57) (*SI Appendix, Supplementary Methods*). Each reaction was performed in triplicate, with the matched non-reverse-transcribed sample run simultaneously. cDNA abundance was calculated using a standard curve and normalized to the reference gene *ALG9.* Oligonucleotides used for qPCR are listed in *SI Appendix, Dataset S2*.

### Chromatin Immunoprecipitation, ChIP-qPCR, and Library preparation

For ChIP-seq experiments (figures 1-3), cells were grown in YPD overnight in 5mL cultures then back-diluted to a concentration of OD600 ∼ 0.1 in 50mL YPD the following day. Cells were grown to mid-log phase (OD600 ∼ 0.6-1.0) and ∼5×10^8^ cells were crosslinked in a final concentration of 2% formaldehyde at room temperature for 15 min. The formaldehyde was quenched using a final concentration of 1.5M of Tris for 5 min.

For Anchor Away ChIP-seq experiments (figures 4,5), cells were grown overnight in YPD, then back-diluted to OD600 ∼ 0.1 in 50mL YPD the following day, then grown for two-three doublings and collected at OD600 ∼ 0.8. Rapamycin (LC Laboratories) was added to a final concentration of 7.5 µM. Additions of rapamycin were staggered such that all time points were ready at the same OD (∼0.8). Samples were fixed and quenched as above. ∼5×10^8^ cells were collected for each sample. 5% S. *paradoxus* cells by OD were spiked into each *S. cerevisiae* sample and processed according to the chromatin immunoprecipitation protocol.

Cell lysis and chromatin immunoprecipitation was performed as described in (57). Details can be found in *SI Appendix, Supplementary Methods*. For all samples, ∼850 μL soluble chromatin were collected for immunoprecipitation, and 50 μL was reserved for Input. For all ChIP samples, 50 µL DynaBeads Protein G magnetic beads (ThermoFisher Scientific) per sample were equilibrated by washing 5x in FA Lysis buffer. IP for 3xV5-Rap1 was performed using 5μL mouse monoclonal V5 (ThermoFisher Scientific). For all 3xFLAG-tagged proteins (Taf1, Sua7, Elf1, Rpb3), IP was performed using 5μL mouse monoclonal anti-FLAG® M2 antibody (Millipore Sigma). Samples were eluted by adding 100 µL TE + 1% SDS to the beads. Input samples were brought to a total volume of 100 µL with TE + 1% SDS. The beads and elution buffer were incubated at 65°C overnight to reverse crosslinking, followed by treatment with RNaseA and Proteinase K. DNA was purified using a QIAquick PCR purification kit (Qiagen).

For ChIP-qPCR, ChIP samples were diluted 10-fold and Input samples were diluted 100-fold in nuclease-free water. Reactions were set up in triplicate and run using the same reagents and parameters as for RT-qPCR above. Abundance was calculated using a standard curve for each primer set, and the ratio of IP/Input was plotted. Oligonucleotides used for this experiment can be found in *SI Appendix, Table S2*.

Libraries were prepared for high-throughput sequencing according to manufacturer’s recommendations using the Ultra II DNA Library Prep kit (NEB). Samples were multiplexed and paired-end sequencing was performed using either a MiniSeq or NovaSeq 6000 (Illumina).

### Alignment and mapping

Sequencing reads were aligned using Bowtie2, using options = “--local --soft-clipped-unmapped-tlen --no-unal --no-mixed --no-discordant” (91) to a reference genome. For standard ChIP-seq experiments (figures 1-3) the genome file was derived from SacCer3 and modified to include, where appropriate, the mutant *HML-p* rap1 binding site mutation, *hml 2::Cre, mat*Δ, and *hmr*Δ, or *hml*Δ *hmr*Δ in the case of *MAT* strains. Analysis was performed using custom Python scripts derived from Goodnight and Rine 2020. Fragments ranging from 0-500bp were mapped, Reads were normalized to the non-heterochromatic genome-wide median (i.e., to the genome-wide median excluding rDNA, subtelomeric regions, and all of chromosome III), and converted to bedgraphs for display. For coverage calculations in figure S2A, peak summits were defined by MACS3 callpeak, using a cutoff of q < 0.01. The positional information and coverage for these peaks can be found in *SI Appendix, Dataset S3*.

For Anchor-Away experiments, a custom, concatenated hybrid genome was generated using modified SacCer3 (unique *HML* sequence, *hmr*Δ) and the *S. paradoxus* genome CBS432 (genbank). Reads were aligned as above. *S. paradoxus* read count served as the normalization factor for each sample. Normalization data be found in *SI Appendix, Dataset S4*.

All displays of ChIP-seq normalized coverage over a defined region were displayed using a custom Rscript and ggplot2.

### Peak-calling and filtering for Anchor-Away experiments

We followed the framework for peak calling and filtering laid out in (56). MACS peak filtering was performed to identify regions of distinct peaks across the *S. cerevisiae* genome in control samples (DMSO-IP, time 0) using a no-tag control sample as the input over which the program defined peaks. Summits were defined using the callpeak function and options “-f BAMPE -g 1.2e7 -q 0.01 --keep-dup=auto -B --call-summits”, identifying 1118 Rap1-bound regions genome-wide. Peaks were defined as 150bp on either side of the summit as defined by MACS. We counted read coverage over each region in duplicate Rap1-depletion sample, and these values were normalized to the *S. paradoxus* read counts per sample as described above. A table containing positional information for these peaks, and the normalized count data, can be found in *SI Appendix, Dataset S5*.

Peaks at each locus were fit using the exponential decay model described in de Jonge et al, filtering for peaks with p-value log(k_off_) < 0.05. The fits were done in R with the nls function using the formula: “nls(ChIP ∼SSasymp(time, yf, y0, log_koff)” (see *SI Appendix, Supplementary Methods*). The 377 peaks used in the analyses for figures 5, S3, and S4 represent peaks that fit the non-linear regression model and were within 300bp upstream of an ORF and/or located in the subtelomeric region (defined as 15kb from the ends of chromosomes). A table containing the calculated fit of the decay curves, average residence time, and further classifications can be found in *SI Appendix, Dataset S6*.

### Other datasets

H3 occupancy genome-wide for analysis of the relationship between Rap1 apparent dwell-time or Rap1 enrichment to nucleosome position was downloaded from GEO Accession GSE97290 (64).

Transcript isoforms were defined using a TIF-seq dataset (GEO Accession GSE39128) (60). We then averaged the corresponding NET-seq signal from four biological replicates in GEO Accession GSE159603 (61).

## Supplementary Methods

### Yeast strain construction

C-terminal tags (Rpb3-3xFLAG:KanMX, SUA7-3xFLAG-KanMX, ELF1-3xFLAG:KanMX, TAF1-3xFLAG:KanMX) were generated by amplifying the 3XFLAG::KanMX sequence from pJR2601 (p3FLAG-KanMX; (92)) with primers that included 40 bp of sequence identity with either side of the amplicon, followed by a transformation. The 3xV5-Rap1 N-terminally tagged allele was generated by amplifying the 3xV5 sequence from pJR3191 pFA61-3xV5-NatMX6 with sequence identity to the N-terminal insertion site on either side, then integrated using CRISPR-Cas9 technology as described (87). Rap1 was tagged N-terminally to avoid genetic manipulations of the Rap1 C-terminus, since doing would likely have interrupted interactions with Sir proteins or Rap1 interacting factors and thus obscured our interpretations in the context of silent chromatin. The rap1 binding site mutation was generated by using CRISPR-Cas9-mediated targeting of the *HML-p* sequence coupled with an oligonucleotide extension that incorporated the 2 bp mutation and obliterated the PAM sequence. The mutant allele of *HML* that included synonymous SNPs was used to distinguish sequencing between *MAT*α and *HMLα* in short sequencing reads, utilized in the Anchor Away experiments, was generated by cloning together synthetic DNA gene blocks (Integrated DNA Technologies) and a previously published allele of *HML* (57), and integrated using CRISPR-Cas9 technology. We recapitulated the Rap1 Anchor Away strain first published in (62) by amplifying the 2xV5-FRB sequence from a plasmid and integrating it, at a sequence corresponding to amino acid 134 in the *RAP1* CDS, into the Anchor Away parent strain (gifted from Craig Peterson, originally from (53)) by CRISPR-Cas9. The *S. paradoxus* strain was generated similarly, but transformed into YSP138 (a gift from the Brar/Ünal labs).

### Validation of epitope-tagged strains

Tagged strains were confirmed by PCR and Sanger sequencing, as well as Immuno-blot analysisTo address the possibility that endogenous tagging of the proteins studied impacted viability or silencing, growth-curves for representative strains were conducted over a 24 hour window. We found no difference in growth rates between wild type yeast and any of our endogenously-tagged strains (figure S1A). Furthermore, introduction of epitope tags to representative strains resulted in no silencing defects as measured by RT-qPCR of *HML*α*2*Δ::*Cre* (figure S1B).

### Calculations of apparent loss-of-silencing rate in CRASH strains

Flow cytometry was done on a BD LSR Fortessa using the BD FACSDiva software (BD Biosciences) and FITC and PE-TexasRed filters. Events were analyzed and processed using FlowJo Software (BD Life Sciences) and the flowAI R package (93) as described in Fouet and Rine 2023. Samples were grouped by population; GFP+ RFP+, GFP+ RFP-, GFP- RFP+ and GFP- RFP-. The apparent silencing-loss rate was calculated by quantifying the number of cells that were transitioning from RFP to GFP expression (those expressing both RFP and *GFP*), divided by the sum of all cells still expressing RFP (RFP+ GFP+ and RFP+ GFP-) (48, 87, 90).

### RNA extraction and RT-qPCR

Briefly, at least ∼2×10^7^ cells were grown and collected by centrifugation for each sample. RNA was purified using the RNeasy Mini Kit (Qiagen 74104; Hilden, Germany) according to manufacturer’s instructions, including on-column DNase digestion (Cat No. 79254). RNA was quantified by NanoDrop, and 2mg of RNA was reverse transcribed using SuperScript III reverse transcriptase (Thermo Fisher Scientific catalog number 18080044) and an ‘anchored’ oligo-dT primer. A matched non-reverse-transcribed sample was generated simultaneously. The DyNAmo HS SYBR Green qPCR kit (Thermo Fisher Scientific F410L), including a Uracil-DNA Glycosylase (Thermo Fisher Scientific EN0362) treatment, was used for qPCR and samples were run using an Agilent Mx3000P thermocycler.

### *S. paradoxus* spike-in

A Rap1-V5 tagged strain of *S. paradoxus* was grown in parallel and fixed at the same saturation. As the *S. paradoxus* cells did not contain all other components of the Anchor-Away methodology, notably the *tor1-1* mutation, they were not exposed to rapamycin. After fixation, cells were washed twice in ice-cold TBS.

### Cell lysis and chromatin isolation

Cells were washed twice in ice-cold TBS and twice in ice-cold FA lysis buffer (50 mM HEPES, pH 7.5; 150 mM NaCl, 1 mM EDTA, 1% Triton, 0.1% sodium deoxycholate) + 0.1% SDS + protease inhibitors (cOmplete EDTA-free protease inhibitor cocktail, Sigma-Aldrich 11873580001). Cell pellets were then either flash frozen or lysed. For lysis, cell pellets were resuspended in 800uL FA lysis buffer + 0.1% SDS and ∼500 µL 0.5 mm zirconia/Silica beads (BioSpec Products; Bartlesville, OK) were added. Cells were lysed using a FastPrep-24 5G (MP Biomedicals; Irvine, CA) with 6.0 m/s beating for 20 s followed by 2 min on ice, repeated four times total. Lysate was transferred to a new microcentrifuge tube, and beads were rinsed with 300uL FA lysis buffer + 0.1% SDS and the remaining lysate was transferred to the same microcentrifuge tube. The cell lysate was transferred to 15 mL Bioruptor Pico tubes along with ∼200μL of the corresponding sonication beads (Diagenode C010200031) and sonicated using a Bioruptor Pico (Diagenode B01060010) for 10 cycles of 30 s ON followed by 30 s OFF. After sonication, samples were spun at 4°C for 30 min at 17 k RCF to pellet cellular debris, and ∼900 μL of the chromatin-containing supernatant was saved.

### Chromatin Immunoprecipitation and sample washes following overnight immunoprecipitation

Beads and antibodies were incubated for >3 hours in an end-over-end rotator at 4°C and then rinsed once with 500uL FA Lysis buffer + 0.1% SDS + 0.05% Tween. All immunoprecipitations were performed in an end-over-end rotator at 4°C overnight in the presence of 0.5 mg/mL BSA (NEB B9000S). Each sample was washed in the following manner, with ∼5 minutes washing by incubating on an end-over-end mixer between each step: 2x washes with FA Lysis + 0.1% SDS + 0.05% Tween; 2x washes with Wash Buffer #1 (FA Lysis buffer + 0.25 M NaCl + 0.1% SDS + 0.05% Tween); 2x washes with Wash Buffer #2 (10 mM Tris, pH 8; 0.25 M LiCl; 0.5% NP-40; 0.5% sodium deoxycholate; 1 mM EDTA + 0.1% SDS + 0.05% Tween); and 1x wash with TE + 0.05% Tween.

After overnight incubation at 65°C, 5 uL 10mg/mL RNase A was added to each sample and incubated for 1 hr at 37°C. 10 µL of 800 U/mL Proteinase K (ThermoFisher Scientific, EO0491) was then added to each ample and incubated for 1 additional hour at 65°C. Each sample was washed in the following manner, with ∼5 minutes washing by incubating on an end-over-end mixer between each step: 2x washes with FA Lysis + 0.1% SDS + 0.05% Tween; 2x washes with Wash Buffer #1 (FA Lysis buffer + 0.25 M NaCl + 0.1% SDS + 0.05% Tween); 2x washes with Wash Buffer #2 (10 mM Tris, pH 8; 0.25 M LiCl; 0.5% NP-40; 0.5% sodium deoxycholate; 1 mM EDTA + 0.1% SDS + 0.05% Tween); and 1x wash with TE + 0.05% Tween.

### Peak calling for figure S2A

Peaks were defined as 150bp on either side of the summit. Peaks were ranked by fold-enrichment and we counted read coverage over the 500 most enriched regions using featureCounts (94).

### *S. paradoxus* normalization factor

Number of mapped read segments were calculated using SAMtools idxstats (95), from which number of reads assigned to the *S. paradoxus* genome were recovered. Total number of *S. paradoxus* reads for each sample was collected, excluding the rDNA due to its vast variability in copy number.

### Peak filtering for anchor away experiment

We evaluated goodness of fit by calculating the standard-deviation of the residuals for each peak (figure S4A). Those peaks that did not fit the non-linear regression model were excluded from further analysis (figure S4B,C). We found 227 Rap1 peaks that were centered over tRNA genes or Ty elements (figure S4A). These loci all displayed relatively short apparent residence times and fit the non-linear regression model well (figure S4A,B). Despite this, we excluded them from further analysis as there is no known connection between Rap1 and Pol III transcribed genes, and it is known that highly transcribed loci are often artifacts of hyper-ChIPability (96, 97). The 377 peaks used in the analyses for figures 5, S3, and S4 represent peaks that fit the non-linear regression model and were within 300bp upstream of an ORF and/or located in the subtelomeric region (defined as 15kb from the ends of chromosomes).

### TIF-seq and NET-seq datasets (previously published)

To visualize summary distribution plots of elongating RNA polymerase II signal as they related to Rap1 binding sites and dwell-time, we first defined full transcripts spanning from transcription start sites (TSSs) to poly-A tracts by identifying the most abundant, stable transcript isoforms in a dataset generated by TIF-seq (Transcript IsoForm sequencing; GEO Accession GSE39128) (60). We then averaged the corresponding Native Elongating Transcript sequencing signal (NET-seq) from four biological replicates in GEO Accession GSE159603 (61).

## Supplementary figures

**Figure S1.**
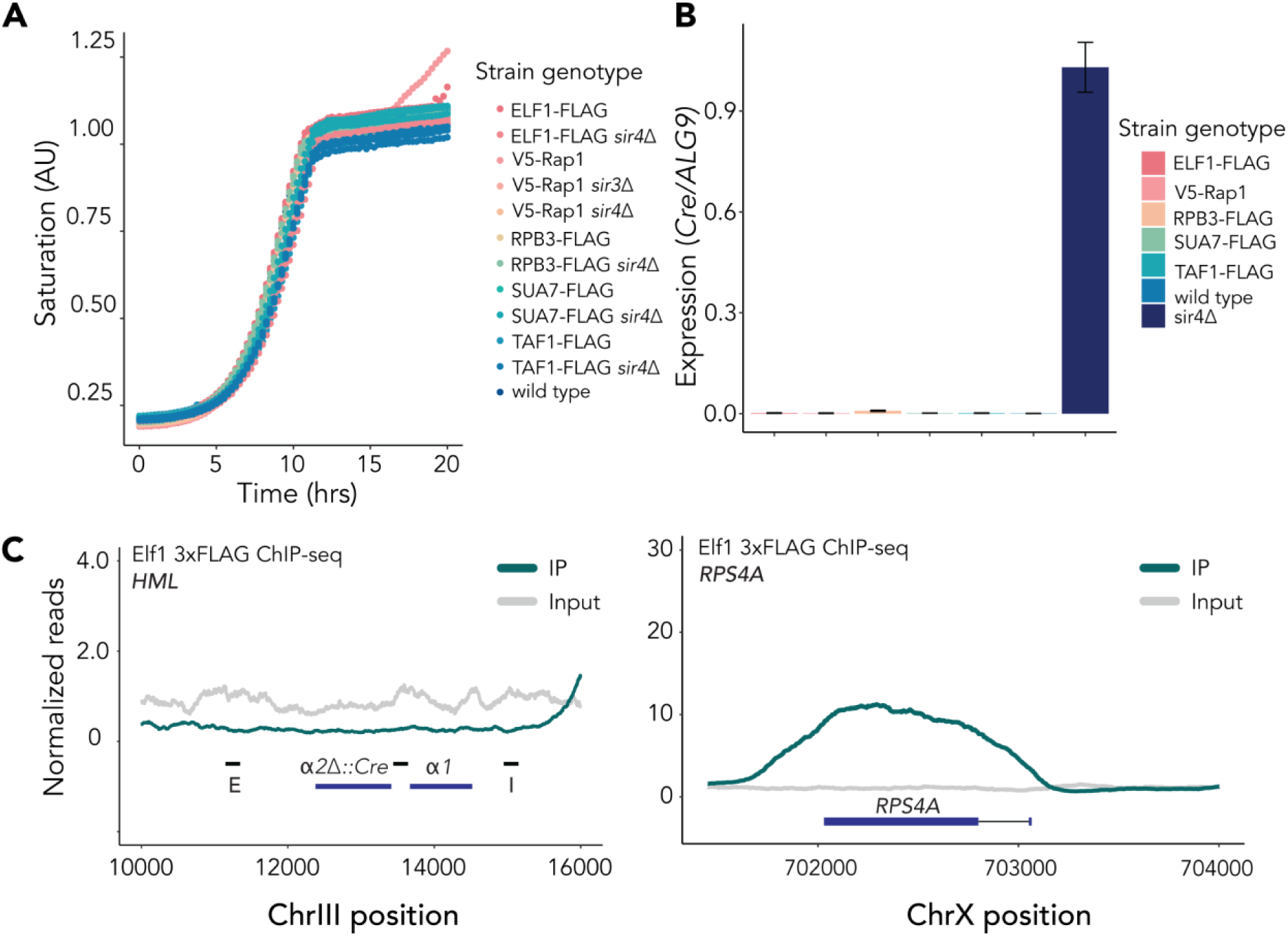
Introduction of epitope tags did not affect viability or silencing. A. Growth curves for representative strains for each of the genotypes listed over 20 hours. B. RT-qPCR quantification of *Cre* expression at *hmlα2*Δ::Cre in representative strains for each of the genotypes listed, normalized to the control locus *ALG9.* Error bars ± SD. C. Averaged normalized reads for ChIP-seq of two Taf1-3xFLAG samples at *HML* (left) and *RPS4A* (right) in *SIR* cells. Black bars represent 200 bp surrounding Rap1 binding sites at *HML-E, HML-p* and *HML-I,* respectively. IP samples are shown in dark green, input values are in grey. Coverage for only one sample is plotted for input. IP and input are plotted on the same scale.

**Figure S2.**
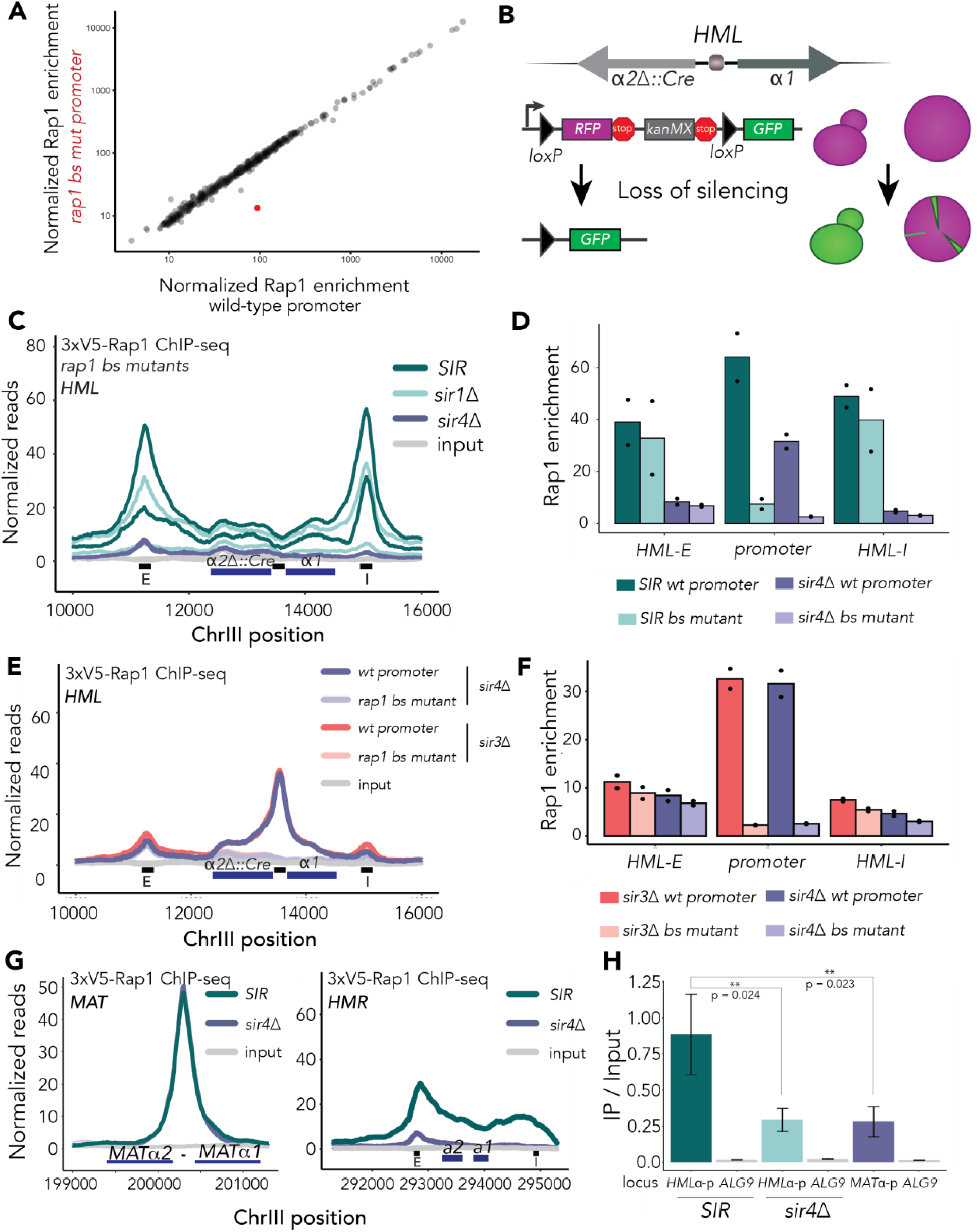
Supporting information for Rap1’s role in strengthening silencing A. Normalized Rap1 ChIP signal for top 500 peaks as defined by MACS in cells with or without the rap1 bs mutation at *HMLp*. Plotted values represent the average of two biological replicates. The peak corresponding to *HMLp* is shaded in red and is the most significantly different between the two. B. CRASH assay experimental design. In this strain, *HMLα2* is replaced with Cre recombinase (*hmlα2*Δ::Cre). A cassette in which an *RFP* and a selectable marker are flanked by loxP sites and driven by the strong *TDH3* promoter resides at the unrelated *URA3* locus. Downstream of loxP-RFP-kanMX-loxP is GFP with no promoter. Upon loss of silencing at *HML*, *Cre* expression induces an irreversible switch from RFP-kanMX expression to GFP expression (supp fig 2B). In a colony, GFP-positive sectors represent a loss-of-silencing event that was enough to allow expression of even one Cre transcript in the cell at the vertex of the sector, with the subsequent progeny represented in the growth outwards. C. Normalized reads mapped to *HML* in two 3xV5-Rap1 in *sir1*Δ (light green), *sir4*Δ (purple) and *SIR* (dark green) ChIP-seq experiments. All strains harbor the two base-pair change in the Rap1 binding motif at the promoter (rap1 bs mut). Light grey represents input samples. Black bars along x-axis represent 200 bp surrounding Rap1 binding sites at *HML-E, HML-p,* and *HML-I,* respectively. IP and input values are plotted on the same scale. D. Quantification of Rap1 enrichment from figure 2A and 2E over each of the three black bars representing *HML-E, HML-I,* and *HML-P.* Black dots represent individual replicates with the mean shown as colored bars. E. Normalized reads mapped to *HML* in 3xV5-Rap1 ChIP-seq experiments. *sir4*Δ wild-type *HMLp* and rap1 bs mutant *HMLp* cells are in dark and light purple, respectively. *sir3*Δ wild-type *HMLp* and rap1 bs mutant *HMLp* cells are in dark and light pink, respectively. Light grey represents input samples. Each plot is the average of two biological replicates. Light grey represents input samples. Black bars along x-axis represent 200 bp surrounding Rap1 binding sites at *HML-E, HML-p,* and *HML-I,* respectively. IP and input values are plotted on the same scale. F. Quantification of Rap1 enrichment from (E) over each of the three black bars representing *HML-E, HML-I,* and *HML-P.* Black dots represent individual replicates with the mean shown as colored bars. G. (Left) Normalized reads mapped to *MAT* in two 3xV5-Rap1 ChIP-seq experiments, averaged. (Right) Normalized reads mapped to *HMR* in two 3xV5-Rap1 ChIP-seq experiments, averaged. Dark green lines represent *SIR* cells. Dark purple lines represent *sir4*Δ cells. Grey lines represent input samples. The Rap1 binding site at the bidirectional promoter is represented by a black line on the x-axis (left). Black bars along x-axis represent 200 bp surrounding *HMR-E, HMR-p,* and *HMR-I,* respectively IP and input values are plotted on the same scale. H. ChIP-qPCR of 3xV5-Rap1 IP / Input enrichment at the *HML*α-promoter in *SIR* and *sir4*Δ, and at the *MAT*α promoter, and each at a negative control locus *ALG9.* N = 3; Unpaired t-test p = 0.024 between *HML*α-promoter in *SIR* and *sir4*Δ; p = 0.023 between *HML*α-promoter and *MAT*α promoter.

**Figure S3.**
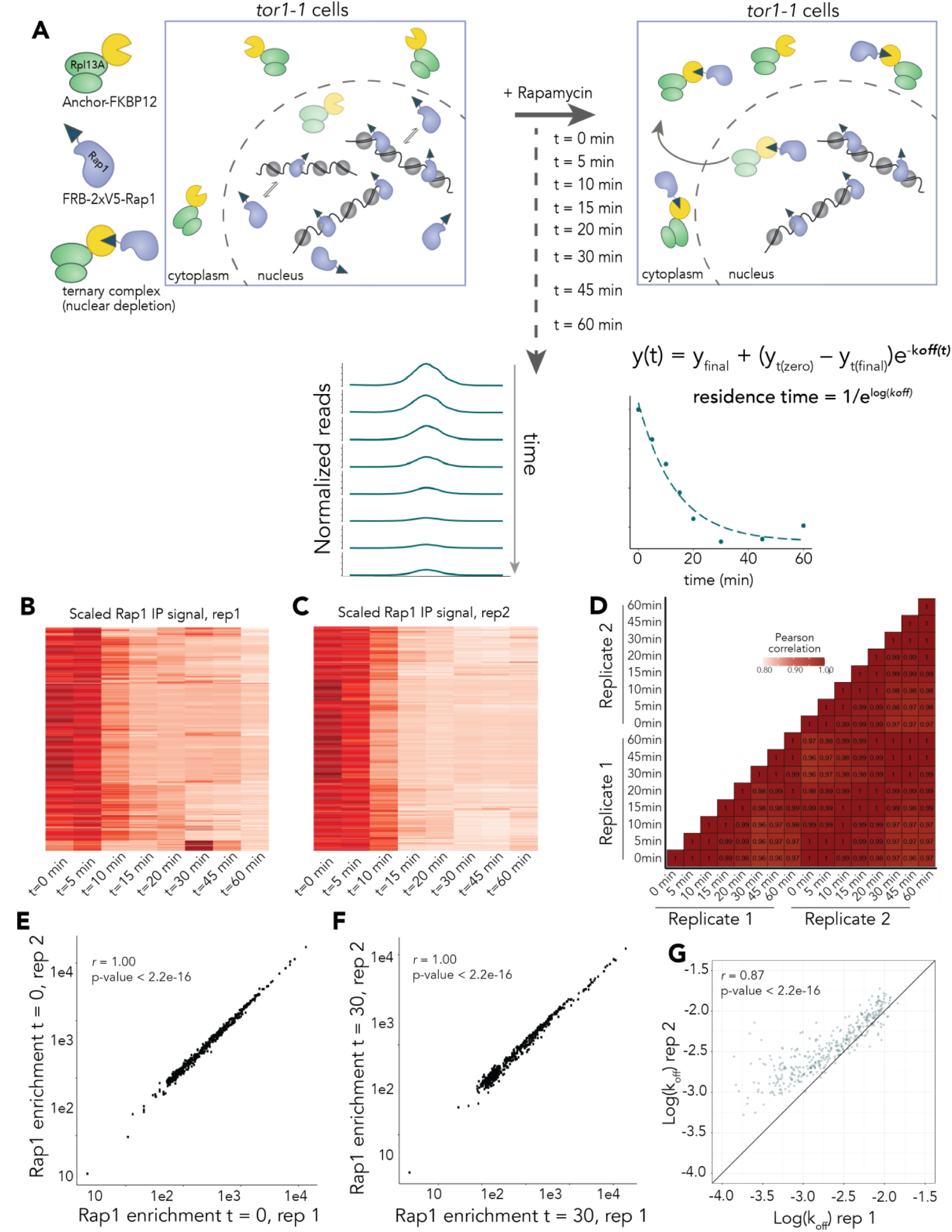
Design and validation of Rap1 anchor away experiment in biological duplicates. A. Experimental setup for Anchor Away. A strain harboring a 2xV5-FRB (FK506 binding protein– rapamycin binding domain) tag at amino-acid 134 in the N-terminus of the protein and the requisite RPL13a-FKBP12 fusion protein, to which rapamycin binds and establishes an interaction surface for the FRB domain, was constructed. Normalized peaks were fit to a non-linear regression model and the k-off rate was extracted. B. Clustered heatmaps of normalized peak coverage over the Anchor Away time-course for biological replicate 1. C. Same as (B) but for biological replicate 2. D. Pearson correlation coefficients, scaled by color, for each pairwise comparison of each time point in two biological replicates. E. Correlation between Rap1 enrichment at time = 0 (DMSO) in replicate 1 on x-axis, and Rap1 enrichment at time = 0 in replicate 2 on y-axis. Pearson correlation *r* = 1.00, p-value < 2.2e- 16. F. Same as (E) but for timepoint t = 30 minutes after addition of rapamycin. Pearson correlation *r* = 1.00, p-value < 2.2e-16. G. Correlation between the calculated log(k_off)_) values for all 377 analyzed peaks for biological replicate 1 on x-axis and replicate 2 on y-axis. Pearson correlation *r* = 0.87.

**Figure S4.**
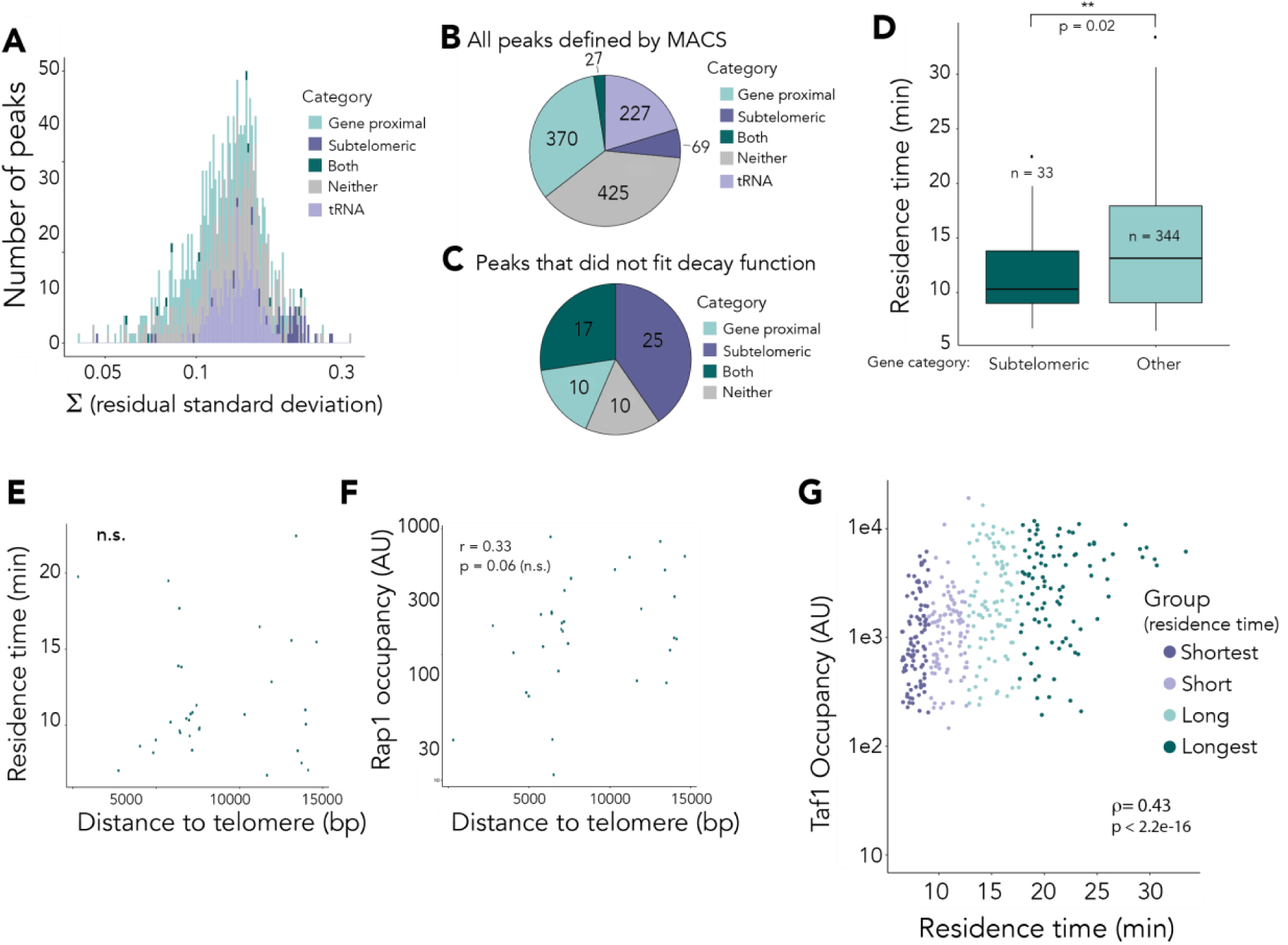
Peak filtering and analysis of subtelomeric Rap1 apparent residence times. All data represent the average of two biological replicates. A. A histogram quantifying the standard-deviation of the residuals for each peak that had a calculated p-value of the log(k_off)_) < 0.05 (n=1056). Peak classifications are denoted by colors in legend. B. A pie chart representing the breakdown of peak classifications of all peaks defined by MACS (see *Methods*) by number (total = 1118). C. A pie chart representing the breakdown of peaks, by classification, from (C) that did not fit the decay function (p-value of the log(k_off)_) > 0.05) (total = 62). D. Box plot quantifying the average residence time for peaks near genes classified as subtelomeric (left, dark green), or all other peaks (right, light green). One-way ANOVA test showing significance p = 0.02. E. Correlation relating distance to telomere in basepairs on the x-axis and average calculated residence time on y axis for peaks categorized as subtelomeric. No significant correlation. F. Correlation relating distance to telomere in basepairs on the x-axis and Rap1 occupancy (enrichment at t=0) on the y-axis for peaks categorized as subtelomeric. Pearson correlation *r* = 0.33, p-value = 0.06 (not significant). G. Correlation between Rap1 apparent residence time (x-axis) and Taf1 enrichment at corresponding Rap1 peaks. Spearman correlation coefficient ρ = 0.43, p-value < 2.2e-16.

## Notes

**Competing Interest Statement:** No competing interests declared.

https://www.ncbi.nlm.nih.gov/geo/query/acc.cgi?acc=GSE227763

### Competing Interest Statement

The authors have declared no competing interest.

### Summary of Updates

This manuscript has been revised to update typos in the abstract and main text.

